# L-tyrosine supplementation is not therapeutic for skeletal muscle dysfunction in zebrafish and mouse models of dominant skeletal muscle α-actin nemaline myopathy

**DOI:** 10.1101/218016

**Authors:** Adriana M. Messineo, Charlotte Gineste, Tamar E. Sztal, Elyshia L. McNamara, Christophe Vilmen, Augustin C. Ogier, Dorothee Hahne, David Bendahan, Nigel G. Laing, Robert J. Bryson-Richardson, Julien Gondin, Kristen J. Nowak

**Affiliations:** Centre for Medical Research, The University of Western Australia, Western Australia, Australia.; Harry Perkins Institute of Medical Research, Western Australia, Australia.; Aix-Marseille University, CNRS, CRMBM, Marseille, France.; School of Biological Sciences, Monash University, Melbourne, Australia.; Centre for Microscopy, Characterisation and Analysis, The University of Western Australia, Western Australia, Australia.; Institut NeuroMyoGène, UMR CNRS 5310 – INSERM U1217, Université Claude Bernard Lyon 1, Villeurbanne, France.; School of Biomedical Sciences, Faculty of Health and Medical Sciences, The University of Western Australia, Australia.; Office of Population Health Genomics, Public Health Division, Western Australian Department of Health, East Perth, Western Australia, Australia.

**Keywords:** nemaline myopathy, L-tyrosine, mouse, zebrafish, muscle, dietary supplementation

## Abstract

Nemaline myopathy (NM) is a skeletal muscle disorder with no curative treatment. Although L-tyrosine administration has been indicated to provide benefit to patients, previous studies have been limited due to sample size or not testing for raised L-tyrosine levels. We evaluated the efficacy of L-tyrosine treatment to improve skeletal muscle function in three animal models of NM caused by skeletal muscle α-actin (*ACTA1*) mutations. Firstly we determined the maximum safest L-tyrosine concentration for inclusion in the water of wildtype zebrafish. We then treated NM Tg*ACTA1*^D286G^-*eGFP* zebrafish from 24 hours post fertilization with the highest safe L-tyrosine dose (10 µM). At 6 days post fertilization, no significant improvement was detected in skeletal muscle function (swimming distance). We also determined the highest safe L-tyrosine dose for dietary L-tyrosine supplementation to wildtype mice. Next we treated the NM Tg*ACTA1*^D286G^ mouse model continuously from preconception with 2% L-tyrosine supplemented to regular feed. We examined skeletal muscles at 6–7 weeks using indicators of skeletal muscle integrity: bodyweight, voluntary running wheel and rotarod performance, all parameters previously shown to be reduced in Tg*ACTA1*^D286G^ mice. The L-tyrosine treatment regime did not result in any improvement of these parameters, despite significant elevation of free L-tyrosine levels in sera (57%) and quadriceps muscle (45%) of treated Tg*ACTA1*^D286G^ mice. Additionally, we assessed the effects of 4 weeks of 2% L-tyrosine dietary supplementation on skeletal muscle function of older (6-7 month old) NM Tg*ACTA1*^D286G^ and KI*Acta1*^H40Y^ mice. This dosing regime did not improve decreased bodyweight, nor the mechanical properties, energy metabolism, or atrophy of skeletal muscles in these NM models. Together these findings demonstrate that with the treatment regimes and doses evaluated, L-tyrosine does not therapeutically modulate dysfunctional skeletal muscles in NM animal models with dominant *ACTA1* mutations. Therefore this study yields important information on aspects of the clinical utility of L-tyrosine for *ACTA1* NM.

**Summary statement:** Despite previous encouraging reports, this study utilising zebrafish and mouse models of nemaline myopathy shows no therapeutic benefit on skeletal muscle functionality in response to L-tyrosine supplementation.

## INTRODUCTION

Tyrosine is a non-essential amino acid that serves as a precursor for several biologically active substances including the brain catecholamine neurotransmitters norepinephrine (NE) and dopamine. Tyrosine may be derived from the diet or via the enzymatic action of phenylalanine hydroxylase on phenylalanine present in the liver, leading to the production of L-tyrosine (the biologically active form of tyrosine; (Deijen et al., 1999). In humans, oral ingestion of L-tyrosine can improve stress-induced cognitive and behavioural deficits (Banderet and Lieberman, 1989; Deijen et al., 1999). Additionally, acute L-tyrosine ingestion is thought to enhance performance via improvements to aerobic power, cognitive performance, neurotransmitter synthesis, and stress related exercise (Luckose et al., 2015). L-tyrosine treatment in rodents can reduce deficits in locomotor activity in old mice following cold water stress, alter stress-related changes in aggression in young mice (Brady et al., 1980), and can protect against both neurochemical and behavioural effects induced by various states of stress (Kabuki et al., 2009; Lehnert et al., 1984).

Dietary supplementation with L-tyrosine may have therapeutic application for patients with the skeletal muscle disorder nemaline myopathy (NM;(Kalita, 1989; Ryan et al., 2008; Wallgren-Pettersson and Laing, 2003). NM is a mainly congenital-onset disorder producing weakened skeletal muscles that contain the characteristic pathological features nemaline bodies (Shy et al., 1963). Twelve different genes can cause NM (Malfatti et al., 2015; Miyatake et al., 2017; Nowak et al., 2015), with a significant proportion of all NM-causing mutations being within the skeletal muscle α-actin gene, *ACTA1* (Nowak et al., 2015). The majority of patients with *ACTA1*-NM have a severe phenotype leading to death within the first year of life (Nowak et al., 2013). At present, no curative treatment exists, highlighting the importance to thoroughly test plausible therapies and for potential novel therapeutic approaches to be identified and investigated.

Daily supplementation of L-tyrosine by an adult male and his 7-year-old son (both with NM) resulted in improved body strength (father), decreased pharyngeal secretions (son), and improved general stamina (both; (Kalita, 1989). After 10 days of L-tyrosine withdrawal, both patients reported reversion to previous clinical conditions, suggesting the improved conditions resulted from L-tyrosine administration (Kalita, 1989). A subsequent small trial contained 5 genetically undefined NM patients (4 infants, 1 adolescent with childhood onset) receiving between 250 to 3000 mg/d of powdered or capsule L-tyrosine for 2 to 5 months (Ryan et al., 2008). Within 72 h on the L-tyrosine regime, all infants displayed initial improvements in “sialorrhoea, skeletal muscle strength and energy levels” (Ryan et al., 2008). Additionally, L-tyrosine (250 mg/d) from 3 months of age was reported to produce marked reduction in oral secretions and improvement in skeletal muscle strength in a severely affected NM patient, however the patient died at 4 months with sudden cardiorespiratory failure (Olukman et al., 2013).

A murine model of NM due to an *Acta1* mutation (KI*Acta1*^H40Y^) was orally dosed via syringe with L-tyrosine (25mg/d) for 4 weeks, from 4 weeks of age (Nguyen et al., 2011). This study concluded that L-tyrosine supplementation alleviated mobility deficits and skeletal muscle pathologies characteristic of KI*Acta1*^H40Y^ mice. However, the study did not address modulatory effects of the L-tyrosine dosing on the early lethality of male mice, nor did it report the sera or tissue levels of L-tyrosine.

Due to the limited, albeit promising data from the few patient studies and the single NM mouse model report, we aimed to comprehensively assess one aspect of the reported therapeutic benefit of dietary supplementation of L-tyrosine, skeletal muscle function. To do so, we chose three dominant NM animal models, each with a missense *ACTA1* mutation resulting in an amino acid substitution (a mouse and a zebrafish model with p.D286G; a mouse model with p.H40Y). In addition to each of these models being suitable animal models of *ACTA1*-NM, they also have characterised deficits in skeletal muscle function ideal for robust assessment of any improvement due to L-tyrosine. Initially, we evaluated different levels of L-tyrosine supplementation in wildtype (WT) zebrafish and mice to identify the highest safe L-tyrosine concentration to dose our NM models. We determined L-tyrosine levels in sera and skeletal muscles of treated mice using this dose, to ensure this supplementation resulted in a significant L-tyrosine increase in the relevant tissues. We tested different L-tyrosine treatment regimes on the dominant *ACTA1*-NM zebrafish and mouse models, and evaluated potential effects on skeletal muscle function using physiological assays and parameters of voluntary exercise.

## RESULTS

### L-tyrosine treatment at higher concentrations can result in deleterious side effects in wildtype zebrafish

A pilot range-finding experiment with WT zebrafish was performed to determine the maximal non-toxic L-tyrosine dose for treatment. Concentrations ranged from 0.1 μM to 10 mM and the survival, heart rate and locomotion of the zebrafish were recorded. Whilst there was a trend between decreasing concentrations of L-tyrosine and resting heart rate, we observed a significant increase in resting heart rate for zebrafish treated with 0.1 μM and 1 μM, suggesting that L-tyrosine is eliciting a biological effect in the fish. The experimental dose for L-tyrosine treatment was determined at 10 μM since zebrafish treated with higher concentrations showed significantly reduced survival and swimming performance compared to water treated controls (**Fig. 1**).

**Fig. 1.**
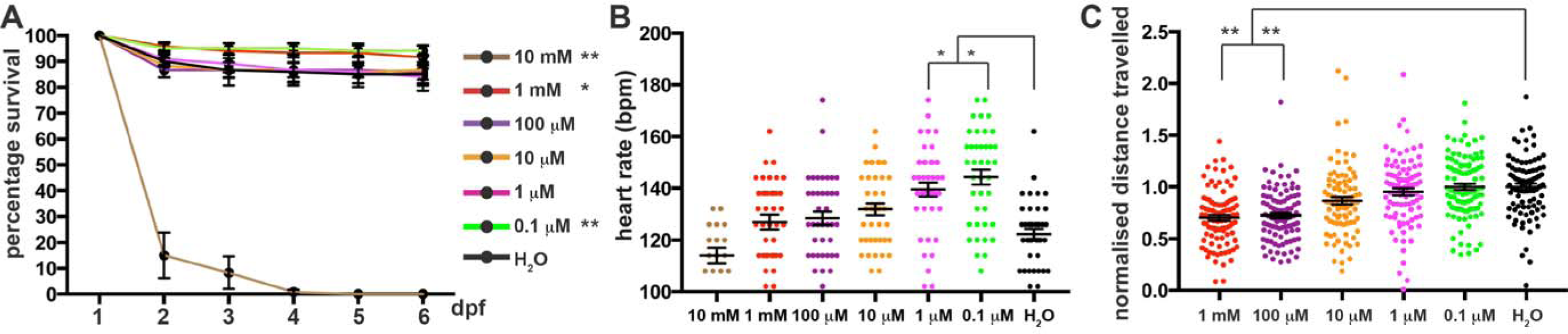
Toxicity analyses for L-tyrosine treatment of wildtype zebrafish. (A) Percentage survival, and (B) resting heart rate in beats per minute (bpm) of zebrafish treated with increasing L-tyrosine concentrations (from 0.1 μM to 10 mM) or H_2_O (used as a control). Error bars represent ±s.e.m. for four independent experiments with 30 zebrafish per experiment for survival assays and 10 zebrafish per experiment for heart rate assays, *p<0.05, **p<0.01 compared to H_2_0 treatment. (C) Normalised distance travelled by 6 dpf zebrafish treated with increasing L-tyrosine concentrations (from 0.1 μM to 1 mM) or H_2_O. Error bars represent ±s.e.m. for four independent experiments with 19-24 zebrafish per dose per experiment. \*\**p* <0.01.

### L-tyrosine addition does not improve the swimming performance of Tg*ACTA1*^D286G^-*eGFP* zebrafish

Tg*ACTA1*^D286G^-*eGFP* zebrafish and their WT siblings (not carrying the ACTA1^D286G^-eGFP cassette) were maintained from 1 day post fertilisation (dpf) in E3 media treated with either 10 μM L-tyrosine or H_2_O until 6 dpf when their swimming performance was assessed. As expected, a significant reduction in distance travelled was observed in water treated Tg*ACTA1*^D286G^-*eGFP* fish compared to control WT siblings (Tg*ACTA1*^D286G^-*eGFP*=0.861±0.021 n=112; WT siblings=1.00±0.073, n=127, *p* <0.01). This deficit in swimming distance in Tg*ACTA1*^D286G^-*eGFP* zebrafish was not ameliorated by the L-tyrosine treatment (water treated=0.861±0.021, n=112; tyrosine treated=0.831±0.082, n=126; **Fig. 2**).

**Fig. 2.**
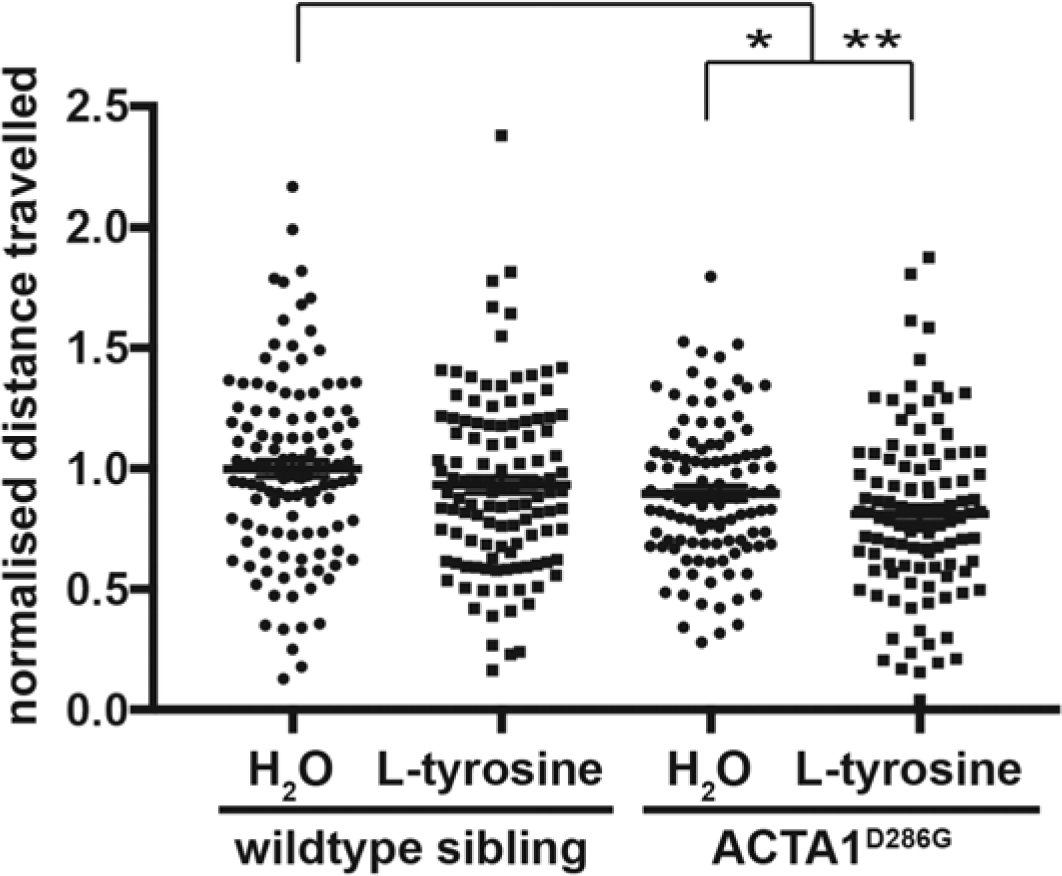
L-tyrosine treatment of ACTA1^D286G^ zebrafish. Normalised distance travelled by 6 dpf Tg*ACTA1*^D286G^-*eGFP* and wildtype sibling control zebrafish treated with 10 μM L-tyrosine or H_2_O. Error bars represent ±s.e.m for five independent experiments with n=238 for Tg*ACTA1*^D286G^-*eGFP*, n=168 wildtype siblings 16-39 zebrafish per treatment, per genotype, per experiment. \**p* <0.05, \*\**p* <0.01.

### Wildtype mice receiving 4% and 8% L-tyrosine supplemented diets after birth display deleterious side effects, whereas mice receiving a 2% L-tyrosine supplemented diet from pre-conception do not

A safety dosing study for L-tyrosine (0, 2, 4 or 8%) supplemented to normal mouse feed (normally 0.7% L-tyrosine) was performed with WT (FVB/NJArc) mice. We observed deleterious outcomes for mice maintained on both the 4% and 8% L-tyrosine supplemented diets. For mice eating the 8% supplemented feed, one dam lost her entire first litter (not necessarily abnormal) then took longer than usual to again become pregnant. She successfully produced a litter of 6 pups, however when the pups were >1 week of age, they were all found dead, a finding out of the ordinary. The second dam receiving the 8% L-tyrosine supplemented feed produced her first and only litter with 2 pups. These pups appeared smaller than usual at the time of wean and were therefore given soft feed located at the base of the cage. Shortly after wean one of the pups was observed to not be moving despite breathing normally and appearing well at an early check that same day. He was therefore sacrificed. Both pups were determined to be ~50% of the weight of WT mice fed the standard diet.

A total of 30 pups were born to the dams receiving the 4% L-tyrosine supplemented feed. Many of these pups, and their mothers, appeared generally dishevelled with ruffled fur. Most pups were found missing on postnatal day 17 (presumably died and were then eaten by the dams or their siblings) with only 6 pups surviving beyond this age (80% mortality). The surviving mice had decreased bodyweight compared to age-matched mice on the control diet (4% L=tyrosine, 11.5±1.7g, n=6; control diet, 15.4±1.4g, n=9, *p* =0.0004).

Due to the animal welfare concerns surrounding these findings, the 4% and 8% L-tyrosine supplemented diets were not further pursued. No detrimental side effects were overtly noticeable for the dams with the 2% L-tyrosine supplemented diet or their resulting offspring (n=17), with all pups surviving beyond wean age and appearing by eye to be similar to those born to mice on the regular diet. Therefore this dose was subsequently evaluated for therapeutic benefit in the two NM mouse models, with the dosing regime being either from pre-conception or commencing at 5 to 6 months of age. Our measurements of average daily feed consumption in adult NM mice indicated that mice continued to eat the same amount of feed once receiving the L-tyrosine supplementation as there was no change in the weight of feed consumed during the 4 week exposure to the 2% L-tyrosine supplemented diet relative to when mice were receiving the normal diet (~3 g/day and ~4.5 g/day consumed for the KI*Acta1*^H40Y^ and the Tg*ACTA1*^D286G^ mice respectively).

### Normal feed supplemented with 2% L-tyrosine significantly elevates levels of L-tyrosine in sera and skeletal muscle of Tg*ACTA1*^D286G^ mice

We assayed samples from Tg*ACTA1*^D286G^ mice receiving the 2% L-tyrosine supplemented feed compared to untreated mice on the control diet and determined that the freely detectable levels of L-tyrosine were significantly elevated in both the sera (untreated mice=52.7±7.8 nmol/ml, treated mice=83.0±13.9 nmol/ml, *p* <0.01) and quadriceps femoris muscle (untreated mice = 0.089±0.021 nmol/mg, treated mice = 0.129±0.032 nmol/mg, *p* <0.05; **Fig. 3**).

**Fig. 3.**
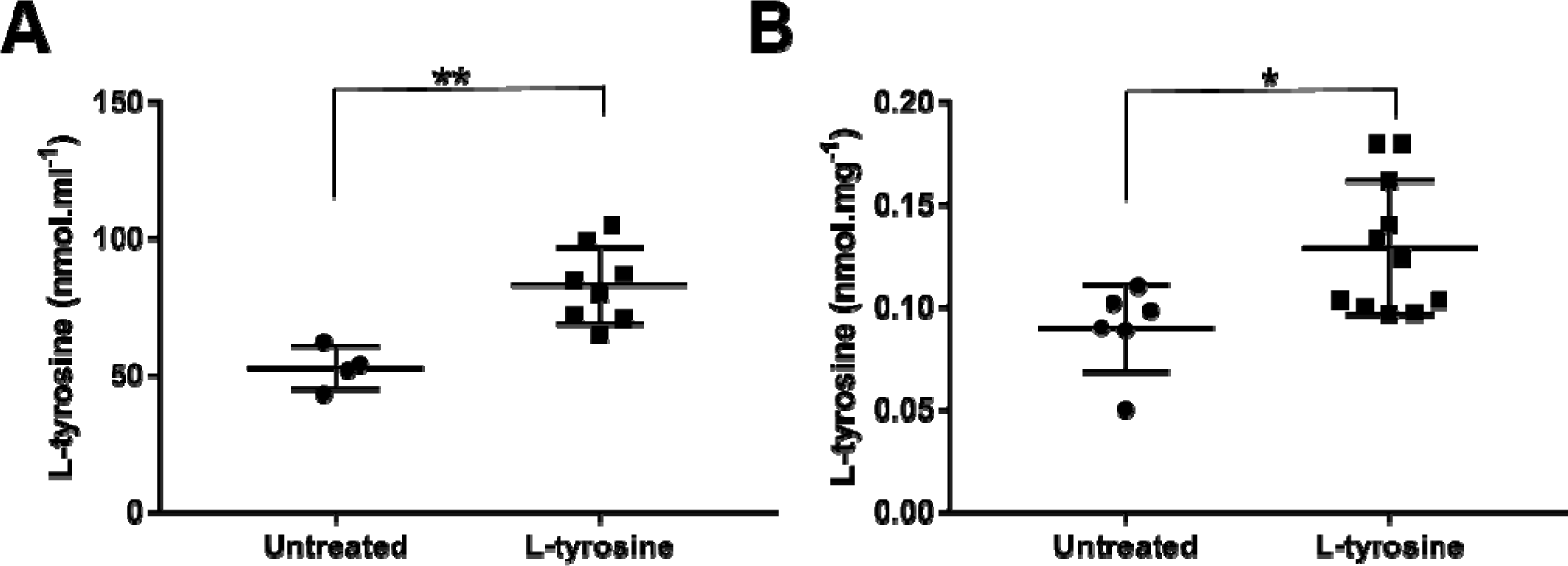
Levels of free L-tyrosine in sera and quadriceps femoris of Tg*ACTA1*^D286G^ mice. The concentration of free L-tyrosine was quantified via LC/MS in (A) sera samples (untreated, n=4; L-tyrosine, n=8), and (B) quadriceps femoris muscles (untreated, n=6; L-tyrosine, n=11) of 6-week old Tg*ACTA1*^D286G^ mice receiving a 2% L-tyrosine supplemented diet from preconception compared to those fed control diets. Each data point represents an individual mouse ±s.d. \**p* <0.05, \*\**p* <0.01.

### Total bodyweight and hindlimb skeletal muscle volume are not increased in L-tyrosine treated *ACTA1*-NM mice

At 6 weeks of age there was no improvement in overall bodyweight of male or female Tg*ACTA1*^D286G^ mice treated from pre-conception (**Fig. 4A**). To the contrary, a significant decrease in total bodyweight was detected in L-tyrosine treated male mice compared to untreated Tg*ACTA1*^D286G^ mice (**Fig. 4A**). Likewise, reduced total bodyweight was not negated at 6-7 months for *ACTA1-*NM mice from either mouse model treated for 1 month (Tg*ACTA1*^D286G^ treated=32.8±4.4g; untreated=34.6±2.7g and KI*Acta1*^H40Y^ treated=22.6±1.7g; untreated=23.8±2.4g; **Fig. 4B**). Additionally, at this older age the hindlimb muscle volume was not different between treated and untreated mice for both models (**Fig. 4C**).

**Fig. 4.**
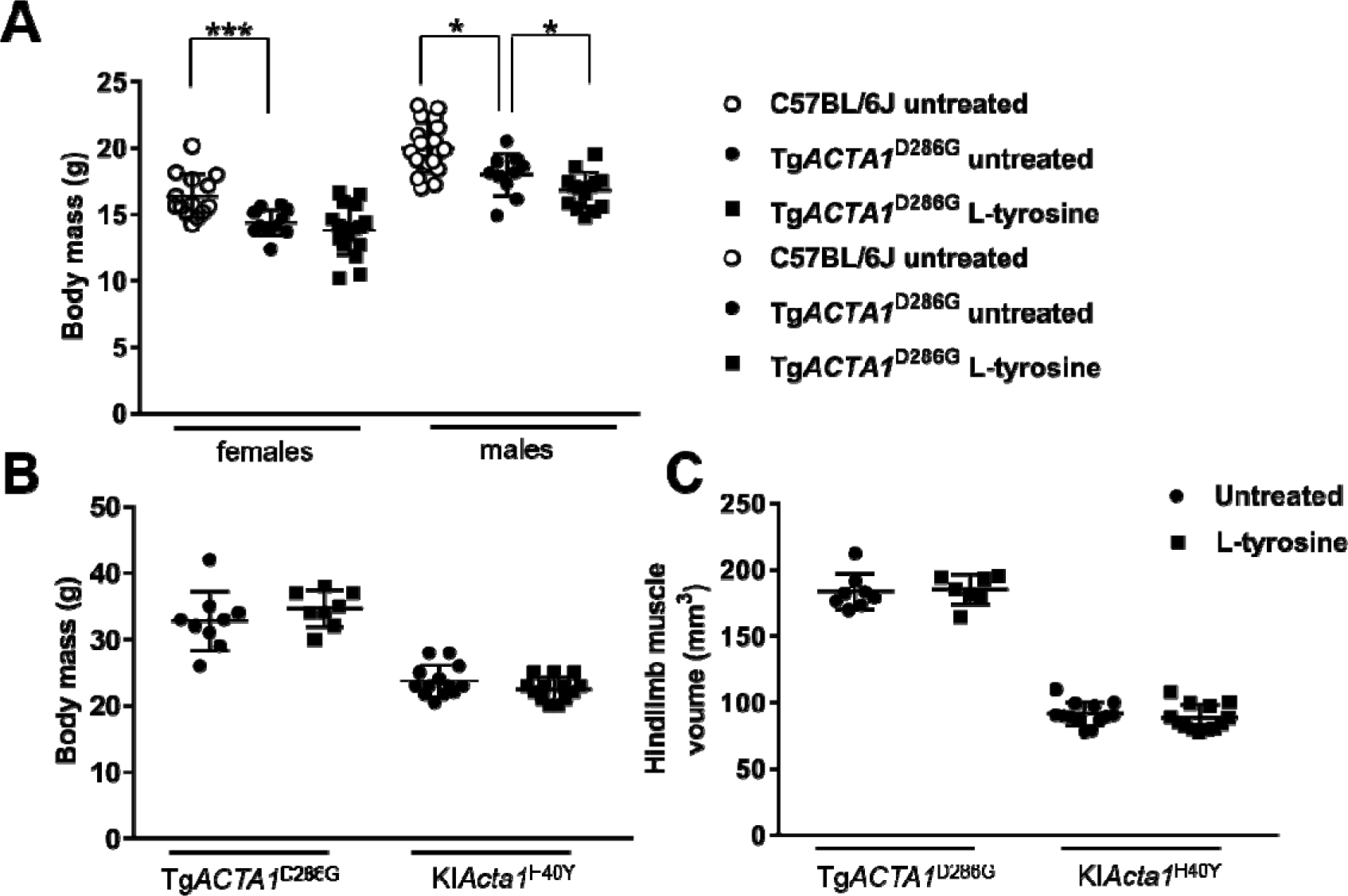
Total body mass and hindlimb muscles volume for NM mice treated with a 2 % L-tyrosine supplemented diet. Total body mass was determined in 6-week old Tg*ACTA1*^D286G^ mice (A) (untreated = 10 males, 12 females; L-tyrosine = 14 males, 17 females) receiving a 2 % L-tyrosine supplemented diet or an untreated diet, as well as C57BL/6J mice (untreated=10 males, 5 females). (B) Total body mass of 6–7 month old mice. Tg*ACTA1*^D286G^ (untreated=9 males, L-tyrosine=8 males,); KI*Acta1*^H40Y^ (untreated=13 females, L-tyrosine=13 females,). (C) Hindlimb muscles volume for mice: male 6–7 month old Tg*ACTA1*^D286G^ (untreated=8; L-tyrosine=7), female KI*Acta1*^H40Y^ (untreated=13; L-tyrosine=13). Each data point represents an individual mouse ±s.d. \**p* <0.05, \*\*\**p* <0.001.

### L-tyrosine treatment does not improve the voluntary running wheel or rotarod performance of Tg*ACTA1*^D286G^ mice

As per Ravenscroft *et al.,* (2011), the voluntary running wheel and rotarod performances of Tg*ACTA1*^D286G^ mice are impaired compared to WT mice. Tg*ACTA1*^D286G^ mice treated from prior to conception did not exhibit any significant improvement for any voluntary running wheel activity parameters relative to untreated mice of the same sex (**Fig. 5**). Similarly, none of the rotarod measurements were significantly improved for treated versus untreated male Tg*ACTA1*^D286G^ mice (**Fig. 6**).

**Fig. 5.**
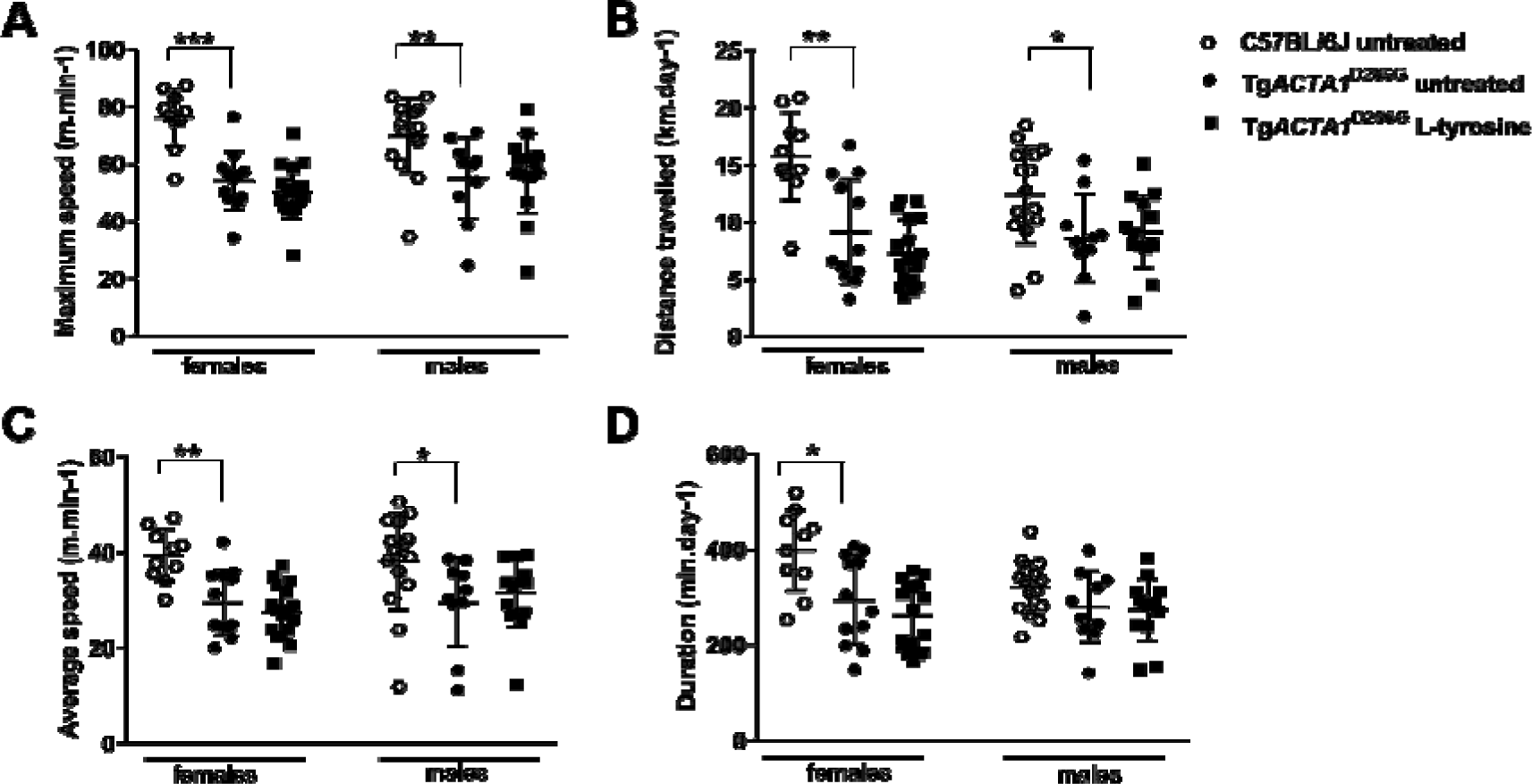
Voluntary running wheel activity in 6-week old Tg*ACTA1*^D286G^ mice receiving a 2% L-tyrosine supplemented diet from preconception compared to Tg*ACTA1*^D286G^ and WT C57BL/6J mice fed an untreated diet. Parameters of voluntary running wheel performance measured included (A) maximum speed (m/min^)^, (B) distance travelled (km/d), (C) average speed (m/min), and (D) duration spent running (min/d). Untreated Tg*ACTA1*^D286G^ mice (10 males, 12 females), L-tyrosine treated Tg*ACTA1*^D286G^ mice (14 males, 17 females), untreated WT C57BL/6J mice (15 males, 10 females). Each data point represents an individual mouse average calculated over days 4, 5, and 6 of voluntary wheel access ±s.d. \**p* <0.05, \*\**p* <0.01, \*\*\**p* <0.001.

**Fig. 6.**
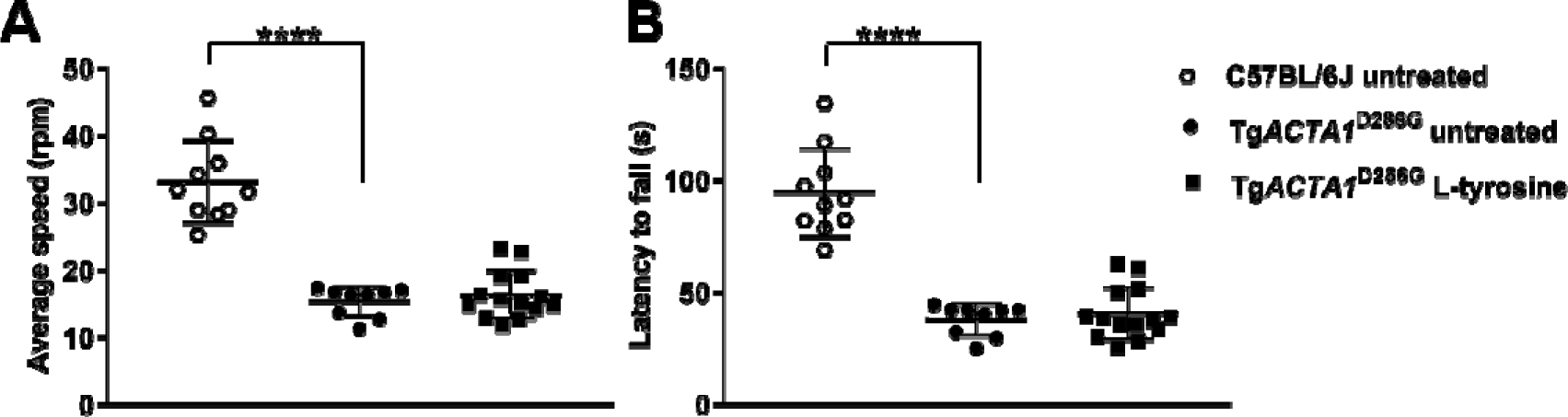
Accelerated rotarod performance in Tg*ACTA1*^D286G^ and C57BL/6J male mice. Performance on an accelerated rotarod apparatus was determined in 6-week old Tg*ACTA1*^D286G^ male mice receiving a 2% L-tyrosine supplemented diet (n=14) compared to Tg*ACTA1*^D286G^ (n=9) and C57BL/6J (n=10) mice fed on untreated diets. (A) Average speed at fall (rpm), and (B) latency to fall (s). Each data point represents the average of 3 attempts by an individual mouse ±s.d. \*\*\*\**p* <0.0001.

### Mechanical performance and metabolism of skeletal muscles in Tg*ACTA1*^D286G^ and KI*Acta1*^H40Y^ mice is not increased by L-tyrosine treatment

Absolute maximal force at 6-7 months of age was unchanged for the two NM mouse models after 1 month of treatment (**Fig. 7A & 7B**). Force production during the fatiguing protocol was comparable for the treated and untreated mice for each model (**Fig. 7C & 7D**). Consequently, the fatigue index (Tg*ACTA1*^D286G^ treated=0.22±0.07; untreated=0.22±0.08 and KI*Acta1*^H40Y^ treated=0.35±0.13; untreated=0.38±0.11) and measures of resting energy metabolism ([PCr] and pH_i_; **Fig. 8**) were similar for both treatment groups for each model. During exercise, PCr consumption, Pi production (data not shown) and pH variations were also similar between the treated and untreated mice for each model (**Fig. 8A**).

**Fig. 7.**
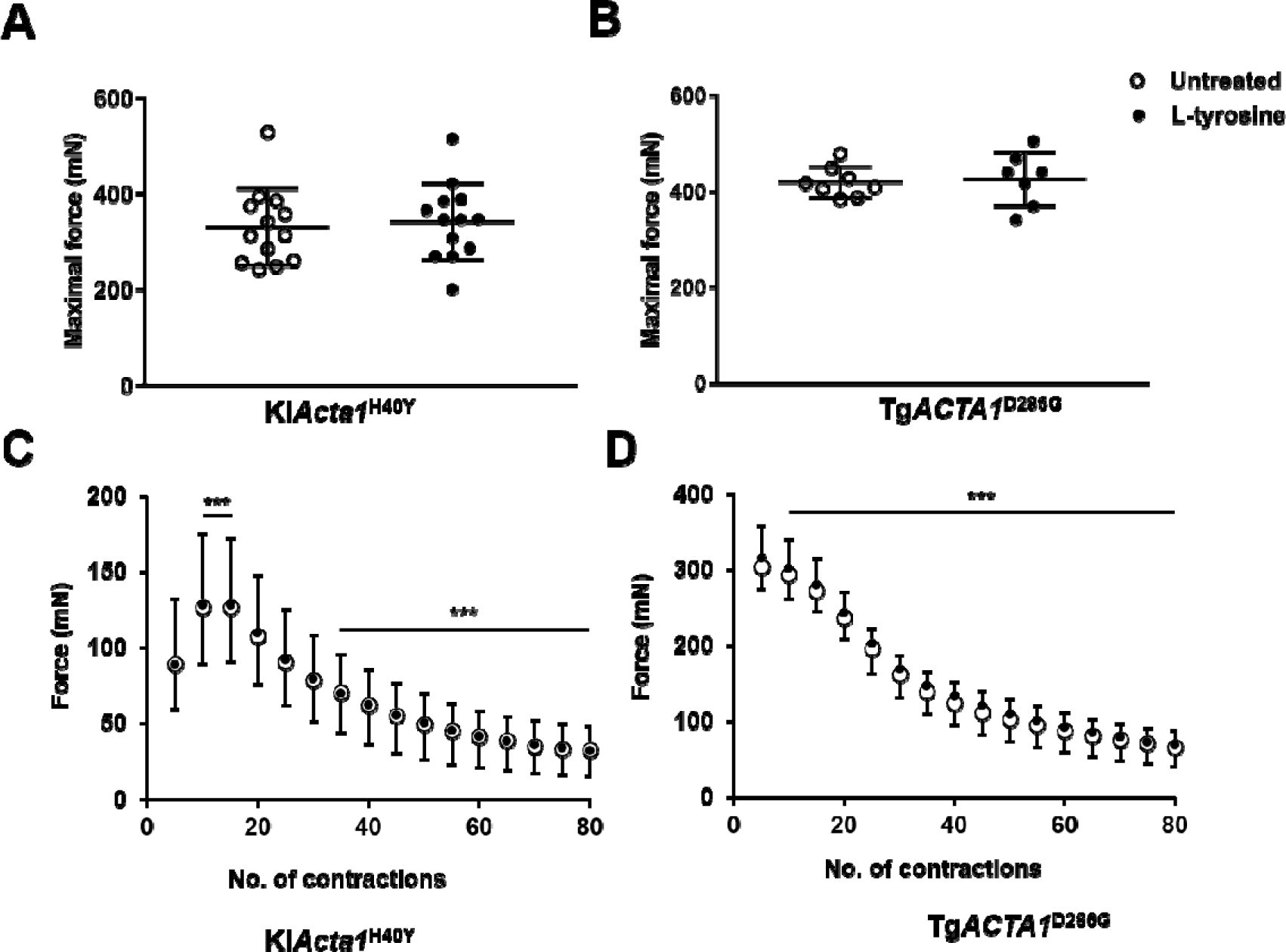
*In vivo* gastrocnemius skeletal muscle performance of Tg*ACTA1*^D286G^ and KI*Acta1*^H40Y^ mice. Absolute maximal force production (A) & (B) and force production during the stimulation protocol (C) & (D) from 6-7 month old Tg*ACTA1*^D286G^ (B) & (D) and KI*Acta1*^H40Y^ (A) & (C) mice fed either an untreated diet or a diet supplemented with 2% L-tyrosine for one month. Tg*ACTA1*^D286G^ (untreated=8 males, L-tyrosine=7 males,); KI*Acta1*^H40Y^ (untreated=13 females, L-tyrosine=13 females). For (A) & (B) each data point represents an individual mouse ±s.d. Data points for parts (C) & (D) are represented by the mean force for 5 contractions of all mice in each diet group ±s.d. \*\*\**p* <0.001 (significantly different from first five contractions), which demonstrates the effect of time on muscle force performance during exercise.

**Fig. 8.**
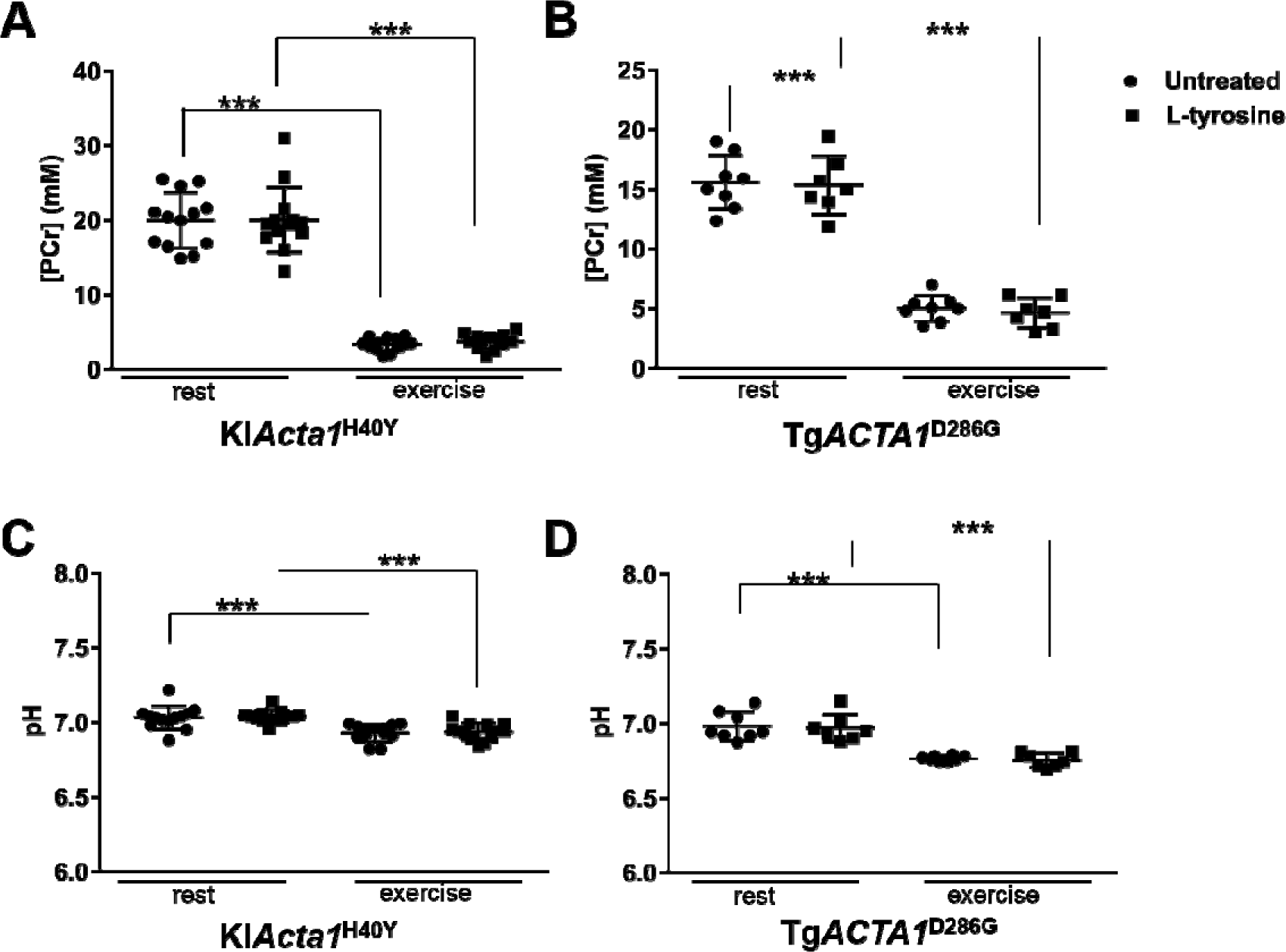
Changes in gastrocnemius PCr and pH during the exercise stimulation in Tg*ACTA1*^D286G^ and KI*Acta1*^H40Y^ mice. [PCr] (A) & (B) and pH (C) & (D) for hindlimb skeletal muscles from 6-7 month old Tg*ACTA1*^D286G^ (B) & (D) and KI*Acta1*^H40Y^ (A) & (C) mice fed either an untreated diet or supplemented with a 2% L-tyrosine diet, at rest versus at the end of the exercise stimulation protocol. The [PCr] and pH values significantly differed between rest and exercised states for each strain, however there was no significant difference detected due to L-tyrosine treatment. Tg*ACTA1*^D286G^ (untreated=8 males, L-tyrosine=7 males); KI*Acta1*^H40Y^ (untreated=13 females, L-tyrosine=13 females). Each data point represents an individual mouse value expressed as mean ±s.d. \*\*\**p* <0.001.

## DISCUSSION

Prior reports of L-tyrosine supplementation to NM patients describe potential positive effects of improved skeletal muscle strength, decreased pharyngeal/oral secretions, and increased stamina/energy levels (Kalita, 1989; Ryan et al., 2008; Olukman et al., 2013), but lacked sufficient numbers for statistical evaluation. The purpose of this study was to evaluate the therapeutic usefulness of L-tyrosine supplementation on one of these previously reported potential benefits, skeletal muscle function, using three dominant *ACTA1*-NM animal models and multiple measures. We utilised one zebrafish (Tg*ACTA1^D286G^-eGFP*) and two mouse (Tg*ACTA1*^D286G^ and KI*ACTA1*^H40Y^) models, encompassing all known laboratory animal models of dominant *ACTA1*-NM. There are a very small number of patients (usually only one individual) with a particular mutation in any of the twelve NM genes, including nebulin and actin, the two most common NM genes. Therefore animal models provide a means to thoroughly investigate a possible therapeutic in multiple individuals with the same genetic composition.

We first investigated the safety of increased L-tyrosine levels for WT zebrafish and determined that higher concentrations of L-tyrosine significantly reduced survival and swimming performance. These findings suggest that the potential toxicity of high L-tyrosine dosing should be considered for humans supplementing with this amino acid, for whatever therapeutic reason. For the zebrafish aspect of our study, we utilised the highest concentration of L-tyrosine that did not produce these negative outcomes (10 µM). Nevertheless, the L-tyrosine treated wildtype and Tg*ACTA1*^D286G^-*eGFP* zebrafish did not exhibit any improvement in the swimming distance travelled.

We also supplemented regular mouse feed with 3 levels of L-tyrosine to determine the highest safe dose. Supplementation with both 4% and 8% L-tyrosine was associated with deleterious side effects in WT mouse mothers as well as their pups, when the feed was supplemented from pre-conception. Our pilot toxicity study in mice was not exhaustive or extensive, yet resulted in high mortality rates for two conditions. These adverse findings with the 4% and 8% doses, especially in combination with the findings from the zebrafish toxicity analysis, provide sufficient reason to highlight possible caution for humans receiving high doses of L-tyrosine.

Tyrosine related toxicity, deleterious effects and weight loss has been previously reported in the literature, e.g. Boctor and Harper, 1968. A potential explanation for the deleterious effects observed in mice receiving higher doses of L-tyrosine may be due to L-tyrosine being a precursor for brain catecholamines. Previous mouse studies report direct correlations between aggressive activity and brain catecholamines in mice (Thurmond and Brown, 1984) with the effects proposed due to the prevention of NE depletion (Deijen et al., 1999). Aggressive behaviour, defined by the number of territorial-induced attacks, was reported in previously unstressed rodents receiving a diet supplemented with 4% L-tyrosine when they were later put under stress (Brady et al., 1980). The authors postulated a reciprocal relationship between dopamine (DA) and NE plus serotonin for the facilitation of aggressive behaviour and suggested that aggressive behaviour may be related to lower brain NE and serotonin levels relative to DA (Brady et al., 1980). Aggression by the mother may have been the cause of death for some of the mouse pups on the 4% and 8% supplemented doses in our study.

As no overtly deleterious side effects were seen with 2% L-tyrosine dietary supplementation, the 2% L-tyrosine supplementation dose was pursued for the efficacy studies with the *ACTA1*-NM mouse models. The 2% L-tyrosine dose significantly increased the free L-tyrosine levels in sera (>55%) and quadriceps muscle (45%) of treated mice. Other studies determined serum tyrosine levels in rats receiving either a 2% or 5% casein diet for 14 days (of 40±3 nmol/ml and 86±8 nmol/ml respectively (Fernstrom and Fernstrom, 1995). The level of sera L-tyrosine detected in untreated Tg*ACTA1*^D286G^ mice (52.7±7.8 nmol/ml) in this study is in accordance with these previous reports. A paucity of data exists for free L-tyrosine levels in rodent skeletal muscles, although baseline levels of L-tyrosine in other tissues (retina, 0.25 nmol/mg; hypothalamus, 0.55 nmol/mg) have been established for rats (Fernstrom and Fernstrom, 1995). The mean value for L-tyrosine in quadriceps muscle of untreated mice we determined (0.089±0.021 nmol.mg^−1^) was less than these levels.

We performed a two-pronged investigation with the 2% L-tyrosine supplemented feed and the NM mouse models, to evaluate pre-birth versus later-onset treatment. We reasoned that if the pre-birth experimental arm established breeding mice on the diet fortified with the highest safe L-tyrosine level and continued the diet through the postnatal and post-wean periods, all offspring conceived would receive the greatest dose and duration of L-tyrosine exposure. Amino acids are known to cross the epithelium of the placental barrier by active transport via specific transporters in syncytiotrophoblast plasma membranes (Jansson, 2001), and are readily detectable in murine breast milk (Rassin et al., 1978). Therefore this L-tyrosine regime would presumably provide the best possible opportunity for prevention/improvement of the skeletal muscle phenotypes attributed to their NM disease if L-tyrosine were therapeutic for this parameter. A well-known example whereby taking supplements prenatally/throughout gestation has significant therapeutic effects is folic acid in the prevention of neural tube defects such as spina bifida (Group, 1991). Tg*ACTA1*^D286G^ mice treated with the pre-birth 2% L-tyrosine supplementation regime until 7 weeks of age demonstrated no improvement in body weight (in fact, L-tyrosine treated 7 week old male Tg*ACTA1*^D286G^ mice weigh significantly less than untreated males), voluntary exercise and rotarod capacity deficits previously reported for this NM model (Ravenscroft et al., 2011).

Our second experimental arm with the murine NM models assessed a dosing regime that started in older mice at 5 to 6 months of age and continued for one month. This is the same treatment duration that Nguyen et al. previously reported for the successful treatment of KI*Acta1*^H40Y^ mice (Nguyen et al., 2011). However, unlike Nguyen et al., we did not detect significant improvements in any phenotype we measured in response to dietary treatment with 2% L-tyrosine for the KI*Acta1*^H40Y^ or the Tg*ACTA1*^D286G^ models. A potential factor that may account for the discrepancies between our findings compared to the previous Nguyen et al. study is the method of delivery and the dose of L-tyrosine. The KI*Acta1*^H40Y^ mice in the Nguyen et al. study were given 25 mg/d of L-tyrosine resuspended in water orally. However as the Nguyen et al. study did not report L-tyrosine levels in sera or skeletal muscles from treated mice we are not able to provide a direct comparison as to the efficiency of the two dosing routes. Thus to relate the dose of L-tyrosine that mice were exposed to in this study to the dose in the previous study, we calculated that mice consuming the 2% L-tyrosine supplemented feed would have received a dose ranging from ~60-90 mg/d (based on the daily intake of 3-4.5 g/d for adult *ACTA1*-NM mice that we determined, which fits with the published murine average daily food consumption range of adolescent mice being from 3.1–6.3 g/d, Bachmanov et al., 2002). Moreover, if we normalise this dose intake to body weight using the average weight for young mice (12g) treated since pre-conception in addition to the older mice (35g) that were treated at 6 months, this equates to 0.5 - 0.75% and 0.17 - 0.25% of total bodyweight for young and older mice respectively. Nguyen et al. reported a similar dose intake of 0.16% of total bodyweight (based on an average weight of 15 g and mice receiving 25 mg/d).

In conclusion, we determined safe concentrations of L-tyrosine for dosing WT zebrafish (water) and mice (dietary supplementation), noting higher concentrations had deleterious effects. The dose evaluated in the dominant *ACTA1*-NM mouse models significantly increased the free L-tyrosine levels in the sera and quadriceps of Tg*ACTA1*^D286G^ mice. Nevertheless, the maximal safe doses utilised had no positive effect on a range of skeletal muscle parameters for TgACTA1^D286G^*-eGFP* zebrafish when added from 24 hpf, nor for the *ACTA1*-NM mice when added continuously from pre-conception (Tg*ACTA1*^D286G^), or for one month from 5 to 6 months of age (Tg*ACTA1*^D286G^ and KI*Acta1*^H40Y^). The amassed data from our multi-pronged evaluation study demonstrate that supplementation of L-tyrosine using the regimes we trialled did not have therapeutic impact on skeletal muscle function in *ACTA1*-NM animal models. Nevertheless, our study does not exclude the potential that L-tyrosine dosing can (i) reduce other symptoms (such as oral secretions and the ability to swallow) that might provide patient benefit, and (ii) have benefit for other genetic causes of NM. However, our findings highlight the imperative to further pursue novel therapies for *ACTA1-*NM.

## Materials and Methods

### Animal ethics statement

All animal experiments were performed in agreement with the guidelines of the respective countries (National Health and Medical Research Council of Australia, French guidelines for animal care, European convention for the protection of vertebrate animals used for experimental purposes, and institutional guidelines). Institutional approval was granted from the respective animal ethics committees (Animal Resources Centre, Monash University, Aix-Marseille University).

### Zebrafish NM model, Tg*ACTA1*^D286G^-*eGFP*)

Zebrafish were maintained according to standard protocols (Westerfield, 2007). Zebrafish strains used were Tg(βAct:loxP**-***mCherry***-**pA**-**loxP:Hs.*ACTA1*^D286G^**-***eGFP*) and Tg(A*ctc1b*:iCre; (Sztal et al., 2015). Crossing of these strains results in the Tg(βAct:loxP-*mCherry*pA-loxP:Hs.*ACTA1*^D286G^-*eGFP*) strain, shortened to Tg*ACTA1^D286G^*-*eGFP*.

### L-tyrosine dosing of the zebrafish NM model

For toxicology testing, 30 WT Tübingen zebrafish were placed in E3 embryo media and treated with increasing doses from 0.1 μM to 10 mM of L-tyrosine disodium salt hydrate (T1145, Sigma, Australia) dissolved in H_2_O. Zebrafish were treated from 24 hpf until 6 dpf. Supplemented embryo media was made fresh and changed daily. Zebrafish were monitored for survival and heart rate as indicators of toxicity. For controls, H_2_O was added to the embryo media instead of L-tyrosine. Four independent treatments were performed for each experiment with 30 fish per treatment.

### Zebrafish swimming assays and resting heart rate determination

The resting heart rates were measured at 2 dpf by counting the number of heart beats in 10 sec. Heart rate measurements were performed in triplicate with 10 fish per experiment. Assay of swimming ability, as well as the subsequent data analyses performed on 6 dpf wild type zebrafish treated with increasing doses of L-tyrosine, Tg*ACTA1*^D286G^-*eGFP* and their control WT siblings treated with 10 µM L-tyrosine were as per (Sztal et al., 2016). Total voluntary distance travelled in a 10-minute period in the dark was measured in mm using zebraboxes (Viewpoint Life Sciences). The values for each genotype and treatment were then normalised to the average of the wildtype untreated siblings in the same replicate. For swimming assays on wild type zebrafish, four independent treatments were performed per experiment with 24 fish per treatment. For swimming assays on Tg*ACTA1*^D286G^-*eGFP* and their control WT sibling, five independent treatments were performed per experiment with 16-39 fish per treatment (238 Tg*ACTA1*^D286G^-*eGFP* fish in total). Based on the pooled SD of the tyrosine and water treated Tg*ACTA1*^D286G^-*eGFP* fish this gave us 0.8 power at 0.05 significance to detect an improvement of 52%. For heart rate and swimming assays all treatments were blinded and randomized to avoid experimental bias. Once the testing and analyses were completed the treatments groups were revealed.

### Mouse NM models Tg*ACTA1*^D286G^ and KI*ACTA1*^H40Y^ and control mouse lines

The Tg*ACTA1*^D286G+/+^ (Ravenscroft et al., 2011); abbreviated to Tg*ACTA1*^D286G^ and KI*Acta1*^H40Y+/-^ (Nguyen et al., 2011); abbreviated to KI*Acta1*^H40Y^ lines were the NM models used in this study. WT mouse strains were used for the initial L-tyrosine safety dosing study (FVB/NJArc), and as a statistical comparison for the Tg*ACTA1*^D286G^ line (C57BL/6J; the closest background strain).

### Dietary L-tyrosine dosing of the NM mouse models

Mouse feed (Speciality Feeds, Glen Forrest, Australia, basal L-tyrosine level of 0.7%; SAFE, Augy, France; basal L-tyrosine level of 0.45%) contained all nutritional dietary parameters either meeting or exceeding the maintenance guidelines for rats and mice outlined by the National Research Council (US; (Animals, 2011). Prior to evaluating the efficacy of L-tyrosine treatment, we conducted a pilot study with normal mice to compare the Australian standard feed (containing 0.7% L-tyrosine) to the same feed supplemented with an additional 2%, 4% or 8% L-tyrosine. Breeding pairs were established on their respective *ad libitum* diets so that all offspring mice were conceived and maintained on these until they were sacrificed at ~7 weeks of age.

Once the 2% L-tyrosine supplemented feed was established as the highest safe concentration of those tested, two dosing regimes were evaluated for their modulation of skeletal muscle disease phenotype. One regime commenced from pre-conception (e.g. dosing of breeding pairs) and continued until sacrifice at 7 weeks of age (Tg*ACTA1*^D286G^ mice: regular ‘untreated’ feed, n=14 males, 17 females; 2% L-tyrosine supplemented ‘treated’ feed, n=10 males, 12 females). The other regime commenced when mice were 5 to 6 months of age and continued for 4 weeks upon which mice were tested (Tg*ACTA1*^D286G^ male mice: treated, n=7; untreated, n=8; KI*Acta1*^H40Y^ female mice: treated, n=13; untreated, n=13). Average weekly feed intake per cage was determined by weighing feed each week for 3 or more weeks prior to addition of the L-tyrosine supplemented feed and then for every week during the 4-week exposure to the treatment. An average daily weight of feed consumed per mouse was then calculated.

### Quantification of L-tyrosine levels in plasma and skeletal muscles of NM mouse models

Blood was collected from L-tyrosine treated and non-treated mice via cardiac puncture at euthanasia at ~7 weeks of age. Immediately afterwards, quadriceps femoris muscles were excised, snap frozen in liquid nitrogen and stored at −80°C. The free L-tyrosine concentration was determined using liquid chromatography/mass spectrometry (LC/MS; University of Western Australia, Centre for Metabolomics, Perth, Australia). Quadriceps femoris samples were thawed from storage at −80°C and weighed prior to being homogenised with ceramic beads in 500 µl of 0.2 M perchloric acid (Hamasu et al., 2009). Sample clean up and derivatization was performed on 100 µl of either muscle lysate or sera using an EZfaast™ Kit (Phenomenex). Then 0.1 µl of sample was applied to an Agilent 1290 UPLC coupled to a 6560 QQQ for the measurement of free L-tyrosine. The EZ:faast AAA-MS 4 μ column 250 × 2.0 mm provided with the kit was used and the acquisition method of the kit was followed. The internal standard was homophenylalanine. Data were acquired in positive ion mode and the transition for L-tyrosine was 396-136.

### Bodyweight of NM mouse models

Male 6-week old Tg*ACTA1*^D286G^ mice treated from pre-conception were weighed prior to individually being housed with access to Low-Profile Wireless Running Wheels (Med Associates Inc, USA) for 6 consecutive days. For both Tg*ACTA1*^D286G^ and KI*Acta1*^H40Y^ mice, body weight was measured after 1 month of exposure to the 2% L-tyrosine supplemented diet or to the normal diet.

### Voluntary running wheel analyses and rotarod assessment of Tg*ACTA1*^D286G^ mice

The daily distance travelled, daily time spent running (summary of 1 min intervals in which at least one wheel revolution was recorded), average speed and maximum speed values were calculated. The mean values for all wheel activity traits on days 4, 5 and 6 were used for data analyses. At ~6 weeks of age, male Tg*ACTA1*^D286G^ mice treated from pre-conception were acclimatised to the rotarod on the day prior to testing by placing mice onto the rotarod at a constant slow speed of 4 rpm for 2 minutes. The following day, mice were tested with placement on the rotarod at 4 rpm, with the rotarod gradually increased speed over 3 minutes until it reached a maximum value of 60 rpm. The test concluded after the mice had fallen off the rotarod. Each mouse was assessed three times on the same day, being allowed at least 10 minutes to rest in between assessments. The latency (time to fall from the rod) and the speed of the rotarod when the mice fell off were recorded for each test. Data were expressed as the averaged value across the three tests.

### Magnetic resonance (MR) and force output measurement in Tg*ACTA1*^D286G^ and KI*Acta1*^H40Y^ mice

MR investigations of Tg*ACTA1*^D286G^ male mice treated for 4 weeks from 6 - 7 months of age were performed in a 4.7-Tesla (T) horizontal superconducting magnet (47/30 Biospec Avance, Bruker, Ettingen, Germany) equipped with a Bruker 120 mm BGA12SL (200 mT/m) gradient insert. Investigations of similarly treated 6 - 7 months of age KI*Acta1*^H40Y^ female mice were performed at 11.75 T on a vertical Bruker Avance 500 MHz.89mm^−1^ wide-bore imager (Bruker, Ettlingen, Germany), equipped with high-performance actively shielded gradients (1 T/m maximum gradient strength, 9 kT.m^−1^.s^−1^ maximum slew rate) and interfaced with Paravision 5.1. A transmit/receive volume RF coil (birdcage, diameter Ø = 3 cm, homogenous length L = 5 cm, Micro 2.5 Probe, Bruker, Ettlingen, Germany) was used for image acquisition.

Mice were anaesthetized and individually placed supine in a home-built cradle specially designed for the strictly non-invasive functional investigation of the left hindlimb muscles. A home-built facemask was incorporated into the cradle and was used to maintain prolonged anesthesia throughout the experiment. The hindlimb was centered inside a ^1^H imaging coil and the belly of the gastrocnemius muscle was located above a ^31^P-magnetic resonance spectroscopy (MRS) surface coil. The foot was positioned on the pedal of the ergometer with a 90° flexion ankle joint. Skeletal muscle contractions were achieved by transcutaneous electrical stimulation using two rod-shaped surface electrodes integrated in the cradle and connected to an electrical stimulator (type 215/T; Hugo Sachs Elektronik-Harvard Apparatus GmbH, March-Hugstetten, Germany). One electrode was placed at the heel level and the other one was located just above the knee joint. Isometric force was measured with a home-built ergometer consisting of a foot pedal coupled to a force transducer. The analog electrical signal from the force transducer was amplified with a home-built amplifier (Operational amplifier AD620; Analog Devices, Norwood, MA, USA), converted to a digital signal, monitored and recorded on a personal computer using the Powerlab 35/series system (AD Instruments, Oxford, United Kingdom).

High-resolution MR images (MRI) were acquired at rest to obtain information about anatomy (i.e. hindlimb muscles volume). For the Tg*ACTA1*^D286G^ mice ten contiguous axial slices (thickness = 1 mm), covering the region from the knee to the ankle, were acquired at rest using a spin echo sequence (echo time (TE) = 18.2 ms; repetition time (TR) = 1000 ms; number of repetition (NEX) = 2; field of view (FOV) = 30 × 30 mm; matrix size = 256 × 256; acquisition time = 8 min 32 s). For the KI*Acta1*^H40Y^ mice fifteen contiguous axial slices (thickness = 0.5 mm), covering the region from the knee to the ankle, were acquired at rest using a gradient echo sequence (TE = 1.5 ms; TR = 189 ms; NEX = 16; FOV = 20 × 20 mm; matrix size = 128 × 128; acquisition time = 6 min 27 s). Images were analyzed with FSLview (FMRIB, Oxford, MS). Regions of interest (ROI) were drawn in the two slices located on the proximal and distal parts of the hindlimb by manually tracing the border of the anatomic cross sectional area of the whole hindlimb muscles. Thereafter, the segmentations of the missing intermediate slices were automatically interpolated (Ogier et al., 2017). The volume of the hindlimb muscles was calculated (mm^3^) as the sum of the volume of the six consecutive largest slices for the Tg*ACTA1*^D286G^ mice or of the nine consecutive largest slices for the KI*Acta1*^H40Y^ mice.

For the measurement of force output, non-invasive transcutaneous electrical stimulation was first elicited with square-wave pulses (0.5 ms duration) on the gastrocnemius muscle of 6-7 month old mice after 4 weeks of dietary treatment. The individual maximal stimulation intensity was determined by progressively increasing the stimulus intensity until there was no further peak twitch force increase. Plantar flexion force was assessed in response to a 100 Hz tetanic stimulation (duration = 0.75 s) and during a fatigue protocol (80 contractions; 40 Hz; 1.5 s on, 6 s off). The peak force of each contraction was measured. Regarding the fatigue protocol, the corresponding tetanic force was averaged every 5 contractions. A fatigue index corresponding to the ratio between the last five and the first five contractions was determined. The resulting force was divided by the volume of the corresponding hindlimb muscles (see above) in order to obtain specific force (in mN.mm^−3^).

### MRS evaluation and metabolic analyses of skeletal muscles from Tg*ACTA1*^D286G^ and KI*Acta1*^H40Y^ mice

Metabolic changes were investigated using ^31^P-MRS at rest and during the fatiguing protocol. Spectra from the gastrocnemius region were continuously acquired at rest and throughout the fatigue protocol. A total of 497 free induction decays (FID) were acquired (TR = 2 s). Data were processed using a proprietary software developed using IDL (Interactive Data Language, Research System, Inc., Boulder, CO, USA). The first 180 FID were acquired at rest and summed together. The next 317 FID were acquired during the stimulation period and summed together. Relative concentrations of high-energy phosphate metabolites (phosphocreatine (PCr), inorganic phosphate (Pi) and ATP) were obtained by a time-domain fitting routine using the AMARES-MRUI Fortran code and appropriate prior knowledge of the ATP multiplets. Intracellular pH (pH) was calculated from the chemical shift of the Pi signal relative to PCr (Moon and Richards, 1973).

### Statistics

Statistical analyses for experiments in Tg*ACTA1*^D286G^ mice and zebrafish were performed using GraphPad Prism 7. All phenotypic traits measured were tested using a nonparametric *t*-test (Mann-Whitney) or a two-way ANOVA. Unequal variances were assumed and all data were tested for normal distribution and passed using D’Agnostino and Perron’s test for Gaussian distribution. All reported sample sizes were powered to detect statistically significant differences in all parameters measured.

Statistical analyses for experiments with 6-7 month-old Tg*ACTA1*^D286G^ and KI*Acta1*^H40Y^ mice were performed with the Statistica software version 9 (StatSoft, Tulsa, OK, USA). Normality was checked using a Kolmogorov-Smirnov test. Two-factor (group x contraction number) analysis of variance (ANOVA) with repeated measures on contraction number were used to compare force production. When a main effect or a significant interaction was found, Tukey’s HSD *post-hoc* analysis was used. One-way ANOVA was used to compare PCr consumption, Pi production and pH_i_. Unpaired *t*-tests were used for body weight, skeletal muscles volume, fatigue index and maximal force comparison. For all mice data shown, values are presented as the mean±standard deviation, with the mean±standard error being reported for all data collected in zebrafish.

## Abbreviations

*ACTA1*: skeletal muscle-actin

*ACTA1-NM*: skeletal muscle-actin nemaline myopathy

ANOVA: analysis of variance

ATP: adenosine triphosphate

CSA: cross-sectional area

DA: dopamine

dpf: days post fertilisation

hpf: hours post fertilisation

FID: free induction decays

LC-MS: liquid chromatography-mass spectrometry

MRS: magnetic resonance spectroscopy

NE: norepinephrine

NM: nemaline myopathy

PCr: phosphocreatine

pH: intracellular pH

Pi: inorganic phosphate

s.d.: standard deviation

s.e.m.: standard error of the mean

T: Tesla

WT: wildtype

## Acknowledgements

We thank the staff at the Animal Resources Centre. Some analysis for this study was performed by the Centre for Microscopy, Characterisation and Analysis (Metabolomics Australia) and was supported by infrastructure funding from the Western Australian State Government in partnership with the Australian Federal Government, through the National Collaborative Research Infrastructure Strategy (NCRIS).

